# Multidimensional isotopic niches inform coexistence mechanisms in an Alpine ungulate community

**DOI:** 10.64898/2026.03.26.714152

**Authors:** Charlotte Vanderlocht, Giorgia Galeotti, Alberto Roncone, Karen Wells, Agostino Tonon, Luca Ziller, Lucrezia Lorenzetti, Matteo Nava, Luca Corlatti, Heidi C. Hauffe, Luca Pedrotti, Francesca Cagnacci, Luana Bontempo

**Author notes:** Francesca Cagnacci and Luana Bontempo equally co-advised this work. Department of Biological, Geological and Environmental Sciences, University of Bologna, Bologna, Italy (Charlotte Vanderlocht).

## Abstract

1. Understanding functional community structure and the niche-based mechanisms that enable coexistence among sympatric species is essential for explaining how biodiversity is maintained in natural systems, and for anticipating how ecological communities will respond to ongoing environmental change. Stable isotope analysis provides a process-oriented perspective on resource use by integrating information across time and space, thereby allowing reconstruction of realised isotopic niches that reflect multiple dimensions of ecological differentiation.
2. We applied this framework to a community of ungulates in the Central-Eastern Italian Alps, including red deer (*Cervus elaphus*), roe deer (*Capreolus capreolus*), and Alpine chamois (*Rupicapra rupicapra*). Using stable isotope ratios in summer-grown hair segments (δ^13^C, δ^15^N, δ^34^S, δ^18^O, δ^2^H), we quantified species-specific n-dimensional niche hypervolumes within a Bayesian framework and estimated niche regions, overlap probabilities, univariate differentiation and multivariate structure.
3. Despite broad dietary overlap typically observed among these ungulates, we found clear isotopic niche segregation, with mean pairwise overlap consistently remaining below 40%. Three dimensions emerged as primary drivers of differentiation: water sourcing (δ^18^O), diet quality (δ^15^N), and habitat openness (δ^13^C). Specifically, chamois appeared to derive more water from plants in their diet rather than from drinking, and to consume a higher-quality diet compared to Cervids. Red deer relied more heavily on forested habitats for resource use compared to roe deer and chamois, and additional isotopic differences between red deer and roe deer may stem from fine-scale abiotic conditions like microclimate and topography. We found no isotopic evidence for differential niche breadth among the three ungulate species.
4. Together, these patterns highlight functional differentiation across multiple ecological axes, offering mechanistic insight into how these ungulates segregate realised niche space despite substantial potential for resource overlap. This multi-element isotope perspective underscores the value of integrative, process-based approaches for understanding current coexistence as well as improving predictions of how mammal communities may reorganise under accelerating environmental change.

## Introduction

Worldwide, ecologically similar species often coexist in natural communities, including phytoplankton (Hutchinson 1961), herbivorous reef fish (Burkepile and Hay 2008), Caribbean lizards (Losos 1994), and African savanna grazers (Prins and Olff 1998). Understanding the mechanisms that enable such coexistence despite overlapping resource use — mechanisms that ultimately underpin much of Earth’s biodiversity — remains a central focus of community ecology (Morin 2009; Abrego and Ovaskainen 2020). In the face of rapid environmental change (Sih et al. 2011) and the growing need to predict community responses with greater accuracy (van der Putten et al. 2010; Ehrlén and Morris 2015; Gaüzère et al. 2018), ecologists are increasingly building on foundational concepts such as the ecological niche (Hutchinson 1957; Holt 2009; Soberón 2014; Laughlin and McGill 2024; Rezvani et al. 2025). These concepts help clarify the mechanisms enabling coexistence today, while also providing a framework for anticipating how community structure and function may shift as species redistribute and local community composition changes (Chen et al. 2011; Lenoir and Svenning 2015; Blowes et al. 2019; Stuart-Smith et al. 2021; Antão et al. 2022).

Multidimensional niche segregation, that is differentiation along one or more niche dimensions (typically dietary, spatial or temporal; Pianka 1973; Schoener 1989; Pocheville 2015) is widely recognised as a key mechanism enabling stable coexistence (Chesson 2000b). However, empirical approaches differ in how they characterise such differentiation. Trait-based metrics (e.g., jaw length, home range size, foraging guild) are frequently used to infer niche differences (McGill et al. 2006; Winemiller et al. 2015; Mouillot et al. 2013; Verberk et al. 2013), yet they rely on assumed relationships between traits and ecological function (D’Andrea and Ostling 2016; Shipley et al. 2016). In contrast, diet- or habitat-based analyses directly quantify resource use (Correa and Winemiller 2014; Kartzinel et al. 2015; Pansu et al. 2022), but typically focus on a snapshot of a single primary niche dimension, while overlooking the others. Moreover, because these approaches often operate on different measurement scales or categorical frameworks, integrating information across dimensions remains challenging (Blonder 2018).

Stable isotope analysis offers a complementary approach that addresses several of these limitations. First, isotopic signatures provide direct measures of assimilated resource use because animals incorporate stable isotopes from their diet, water, and environment into their tissues over time. Second, in sequentially grown and metabolically inert tissues such as keratin, isotopic information is temporally integrated across the period of tissue synthesis (Dalerum and Angerbjörn 2005; Rogers et al. 2020), overcoming the snapshot limitation of conventional diet analyses. Third, isotopes can simultaneously reflect multiple aspects of ecological differentiation. Isotopic ratios of bio-elements such as carbon and nitrogen can inform on primary energy sources (Finlay and Kendall 2007; Lopes et al. 2015), trophic position (Post 2002), habitat openness (Drucker et al. 2010; Krigbaum et al. 2013), and even diet quality (Fantle et al. 1999; Vanderklift and Ponsard 2003; Hobson 2008), while sulphur, hydrogen, and oxygen can capture variation in space use (latitude, elevation, local geochemistry; Ambach et al. 1968; Richards et al. 2003; Bowen et al. 2005) and climatic conditions (Dansgaard 1964; Siegenthaler and Oeschger 1980; Poage and Chamberlain 2001). Last but not least, these signatures are quantified as continuous ratios of heavy to light isotopes (e.g., δ^13^C, δ^15^N, δ^34^S, δ^2^H, δ^18^O), and expressed in identical units (‰): they can therefore be integrated within a common quantitative framework, providing functionally tractable and ecologically meaningful measures that are comparable across species and ecosystems (Bearhop et al. 2004; Layman et al. 2012). Stable isotope analysis thus provides a process-oriented perspective that integrates information across dimensions, enabling reconstruction of realised isotopic niches (Newsome et al. 2007; Boecklen et al. 2011; Newsome et al. 2012) and their mathematical representation as multidimensional hypervolumes (Swanson et al. 2015; Blonder 2018). Such integrative approaches are particularly valuable in ecosystems undergoing rapid ecological reorganisation, where understanding the mechanisms structuring species coexistence is critical for predicting community responses.

Mountain ecosystems are one such context. In the Italian Alps, climate change has altered abiotic conditions and resource dynamics, including a 1.5°C increase in summer temperatures over the past thirty years (Bright Ross et al. 2021), while socio-economic shifts have driven substantial land use change, notably a marked expansion of forest cover (Passoni et al. 2024), as well as changes in wildlife management, including species protection and regulated hunting. Together, these drivers have strongly reshaped the distribution, abundance, and interactions of large mammals (Passoni et al. 2024; Vanderlocht et al. 2026). In this rapidly evolving context, predicting community responses requires identifying the mechanisms that structure species coexistence, in particular, the extent and dimensionality of niche segregation among sympatric species. Here, the Alpine large herbivore guild includes Cervids (red deer *Cervus elaphus*, roe deer *Capreolus capreolus*) and a Bovid (Alpine chamois *Rupicapra rupicapra*). Red deer are large-bodied mixed feeders (Hofmann 1989) and partially migratory (Cagnacci et al. 2024; Galeotti 2024); chamois are also mixed feeders, though less inclined toward roughage consumption (Hofmann 1989) and more specialised to high-alpine habitat (von Elsner-Shack 1985; Mustoni et al. 2005); and roe deer are smaller generalist concentrate selectors (Hofmann 1989) that preferentially use ecotonal habitats (Reimoser and Gossow 1996; Mustoni et al. 2005). Despite these differences, these sympatric ungulates often exhibit broadly overlapping diets (Redjadj et al. 2014; Lioce 2023). Recent population recoveries (Passoni et al. 2024), together with climate and land-use change, are driving upward shifts in summer ranges (Servizio Sviluppo Sostenibile e Aree Protette 2022) and potentially increasing habitat overlap among species. These dynamics underscore the need to reassess spatial, dietary, and functional niche relationships in Alpine ungulates, even among long-coexisting species.

Here, we quantified multidimensional isotopic niches of sympatric red deer, roe deer and chamois populations in the central and eastern Alps, using stable isotope analysis of carbon, nitrogen, sulphur, hydrogen and oxygen (δ^13^C, δ^15^N, δ^34^S, δ^2^H, δ^18^O) measured in summer-grown hair segments. We constructed five-dimensional population-level niche hypervolumes and estimated niche size, overlap and differentiation among species. We assumed broadly comparable incorporation of dietary and water isotopes into hair across species (Vanderlocht et al. *In prep.*).

We hypothesised that sympatric red deer, roe deer and chamois exhibit niche segregation (Chesson 2000b), consistent with classical niche theory (Gause 1934; Hutchinson 1957). We expected significant differentiation along at least one isotopic dimension, with some overlap due to niche filtering (Keddy 1992; Zobel 1997). Given their generalist ecology, larger body size, and larger home ranges, red deer were predicted to have the broadest isotopic niche. We further expected niche differences among the three species to reflect both environmental gradients and variations in realised resource use (Table 1), consistent with niche differentiation.

**Table 1.** Predicted population-level isotopic patterns among sympatric Alpine ungulates, including their ecological interpretation, underlying mechanisms, and key references.

| Stable isotope ratio | Ecological interpretation (ungulate context) | Mechanism (with key references) | Predicted pattern |
| --- | --- | --- | --- |
| $\delta^{13}\text{C}$ | Reflects habitat use for feeding behaviour (in a pure $\text{C}_3$ ecosystem): use of closed habitat leads to lower $\delta^{13}\text{C}$, whereas use of more open areas leads to higher $\delta^{13}\text{C}$. Degree of depletion in $^{13}\text{C}$ is correlated with forest density.<br>*Examples of ungulate studies: Drucker et al. 2008; Drucker and Bocherens 2009; Drucker et al. 2010.* | This phenomenon is known as the ‘canopy effect’ (van der Merwe and Medina 1991; Heaton 1999; Bonafini et al. 2013), and is driven by<br>(1) photosynthetic recycling of $^{13}\text{C}$-depleted $\text{CO}_2$ derived from soil decomposition and/or vegetation respiration (Vogel 1978; Da Silveira et al. 1989) in poorly ventilated forested habitat; and<br>(2) photosynthesis under low light intensity and low $\text{CO}_2$ concentration, involving changes in stomatal conductance and Rubisco activity (with isotopic fractionation during stomatal diffusion and carboxylation; Farquhar et al. 1982; Ehleringer et al. 1986; Broadmeadow et al. 1992).<br>Shade/obscurity, atmospheric $\text{CO}_2$ concentration and $\text{CO}_2$ recycling decrease along closed-to-open habitat gradients. | Red deer and roe deer display lower $\delta^{13}\text{C}$ (using more forested habitats) than chamois (using more high-elevation open areas). |
| $\delta^2\text{H}$ | Reflect elevational occupancy: use of increasingly high elevation leads to decreasing $\delta^2\text{H}$ and $\delta^{18}\text{O}$ (at a given latitude, distance from water masses, and season).<br>*Examples of ungulates studies: Hermes et al. 2017; Morandi et al. 2021.* | This phenomenon, known as the ‘altitude effect’ (Dansgaard 1964; Poage and Chamberlain 2001), is the gradual altitudinal depletion in heavy isotopes of precipitation water resulting from<br>(1) preferential condensation of heavy isotopes, with progressively higher condensation levels and lower condensation temperatures along the cloud’s orographic ascent and cooling (Dansgaard 1964); and<br>(2) preferential evaporation of light isotopes in the falling raindrops (Ambach et al. 1968), especially in summer due to higher temperatures (Siegenthaler and Oeschger 1980).<br>Thus, as an air mass rises orographically, it progressively rains out moisture cooling which becomes increasingly depleted in heavy isotopes | Red deer and roe deer display higher $\delta^2\text{H}$ (occupying lower elevations) than chamois (occupying higher elevations). |
| $\delta^{18}\text{O}$ | | | |
|  |  | (<sup>2</sup>H, <sup>18</sup>O) as condensation temperature decreases under decreasing atmospheric pressure, increasingly so as the air mass itself becomes gradually depleted (Rayleigh condensation, i.e. the process of immediate removal of the condensate — here heavier rain molecules — from the vapour after formation). Additionally, as altitude increases, rainfall distance from the cloud to the ground decreases, leading to decreasing heavy-isotope-enriching evaporation. |  |
| $\delta^{15}\text{N}$ | Reflects feeding strategy and dietary protein content through increased diet-tissue discrimination for more fibrous diets vs more proteic diets, on the grazing-to-browsing continuum.<br>*Examples of ungulate studies: Stewart et al. 2003; Drucker et al. 2010.* | Diet-tissue nitrogen isotope discrimination increases as dietary C:N ratio increases (Cormie and Schwarcz 1996; Fantle et al. 1999; Vanderklift and Ponsard 2003), that is, as the fiber-to-protein content increases or as diet quality decreases (Hobson 2008). As dietary protein content increases, diet-tissue fractionation decreases with the increasing ratio of urine:faecal nitrogen, with urine being <sup>15</sup>N-depleted and faeces being <sup>15</sup>N-enriched (Sponheimer et al. 2003), or in more extreme cases of poor diet quality, the animal must catabolise its own lipids and proteins for energy (Fantle et al. 1999), keeping heavy nitrogen and excreting light NH<sub>3</sub><sup>+</sup> through the urine.<br>Along the browser-grazer continuum of ruminants (Hofmann 1989), browsers (or concentrate selectors) tend to feed on less fibrous and more nitrogen-rich plants than grazers (Hofmann and Stewart 1972; Owen-Smith 1982; McNaughton and Georgiadis 1986). | Red deer and chamois display higher $\delta^{15}\text{N}$ (mixed feeders with higher dietary fiber content) than roe deer (concentrate selectors with higher dietary protein content). |
| $\delta^{34}\text{S}$ | Reflects bedrock and soil composition through limited diet-tissue discrimination.<br>*Example of ungulate study: Kabalika et al. 2020.* | Due to the limited diet-tissue sulphur discrimination throughout trophic levels (Krouse and Herbert 1988), $\delta^{34}\text{S}$ values tend to reflect the local geology (Richards et al. 2003; Stack and Rock 2011). | No difference in $\delta^{34}\text{S}$ expected among chamois, red deer and roe deer, due to population-level overlapping ranges at geological and limnological scales. |

## Material and methods

### Study area and sampling

The study area was located in the Italian Alps (Fig.1), specifically in the Regions of Lombardy and Trentino-Alto Adige/Südtirol, including Stelvio National Park (IUCN category II). Here, rugged mountainous terrain (900-3905 m a.s.l.) features alpine pastures at higher elevations and coniferous forests with managed meadows at lower altitudes. Hair samples (tufts of 5-10 mm diameter of winter coat guard hairs from the thigh of red deer, roe deer and chamois) were collected from hunted individuals or occasional carcasses (e.g. roadkill) between September 2022 and February 2023, in collaboration with hunters and provincial staff involved in the park’s fieldwork and culling programme (Servizio Sviluppo Sostenibile e Aree Protette 2022). Samples were georeferenced and timestamped, transferred at ambient temperature to the Fondazione E. Mach, and stored in dry, dark conditions inside paper envelopes until analysis.

**Figure 1.**
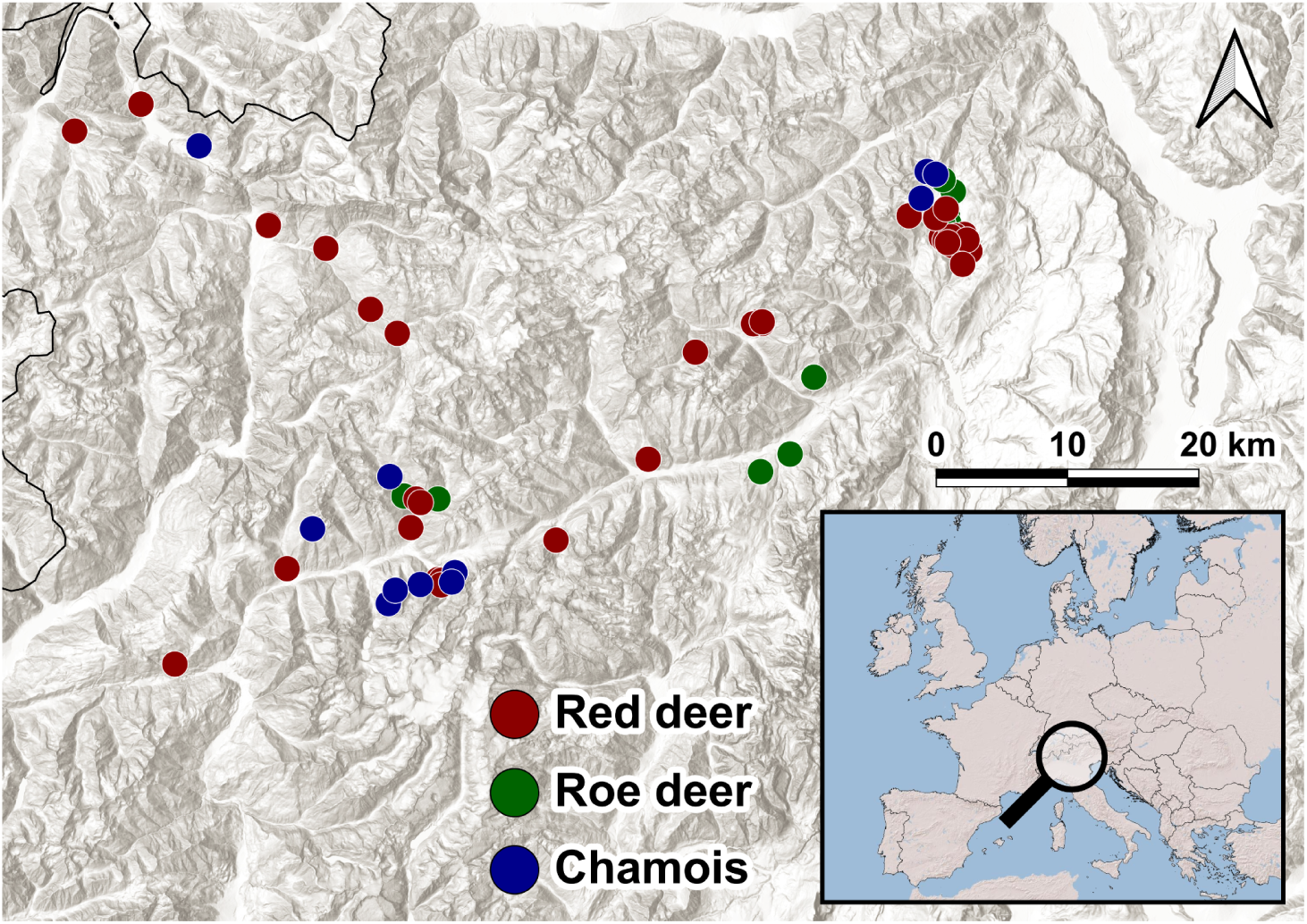
Map with the distribution of ungulate hair samples collected for isotopic analysis. The samples depicted are from red deer (red), roe deer (green), and chamois (blue).

### Sample preparation and isotope analysis

Hair samples were cleaned and defatted via three successive 1-minute sonicated solvent baths (diethyl ether:methanol 2:1; Bontempo et al. 2014). We selected long, intact guard hairs (including root and tip), removed the root to eliminate cellular material and isolate keratin, and segmented each hair into three isometric segments: base (winter growth; Thomas 1997; Mustoni et al. 2005), centre (discarded to reduce seasonal variability), and tip (mid- to late-summer growth; Thomas 1997; Mustoni et al. 2005). In order to reduce temporal variability in growth periods and thereby improve the ecological relevance of comparisons at that scale, only the summer-grown tip segments were retained, weighed (Data S1), and analysed via stable isotope mass spectrometry.

Values of δ^13^C, δ^15^N and δ^34^S were simultaneously determined using a Vario Isotope Cube isotope ratio mass spectrometer (Elemental, Germany), whereas values of δ^2^H and δ^18^O were determined through pyrolysis combustion using a TC/EA (Thermo Finnigan, Germany) interfaced with a Delta Plus XP (Thermo Finnigan, Germany) continuous-flow isotope ratio mass spectrometer (Data S1). Isotope ratios were expressed in δ-notation against VPDB (Vienna Pee Dee Belemnite) for δ^13^C, Air-N2 (atmospheric nitrogen) for δ^15^N, VCDT (Vienna Canyon Diablo Troilite) for δ^34^S, VSMOW (Vienna Standard Mean Ocean Water) for δ^2^H and δ^18^O, and were calculated based on keratin standards and a working in-house standard (wheat) which was calibrated against international reference materials (Data S1) using multi-point normalisation (Brand et al. 2014). Method uncertainty (expressed as one standard deviation) was 0.1‰ for δ^13^C, 0.2‰ for δ^15^N, 0.3‰ for δ^34^S, 0.3‰ for δ^18^O and 2‰ for δ^2^H.

### Replication statement

Although structured at the population level (Table 2), results were used to infer community-level niche structure and overlap among coexisting species.

**Table 2.** Replication statement, indicating the scales at which we sought to make inferences; the scales at which the factor of interest were measured; and the number of replicates for each level of factor.

| Scale of inference | Scale at which the factor of interest is measured | Number of replicates at the appropriate scale | Note |
| --- | --- | --- | --- |
| Species/Population | Individual | Red deer: 50<br>Roe deer: 16<br>Alpine chamois: 29 | One sample per individual, selection of tip segments within hair tuft for temporal resolution |
| Technical replication | Sample (here, tip segments per individual) | 3 technical replicates for $\delta^{13}\text{C}$, $\delta^{15}\text{N}$, $\delta^{34}\text{S}$ per sample | Individual $\delta^{13}\text{C}$, $\delta^{15}\text{N}$, $\delta^{34}\text{S}$ values were calculated as the mean of technical replicates |
| | | 2 technical replicates for $\delta^2\text{H}$, $\delta^{18}\text{O}$ per sample | Individual $\delta^2\text{H}$, $\delta^{18}\text{O}$ values were calculated as the mean of technical replicates |

### Data analysis

Considering a five-dimensional isotopic niche space (δ¹³C, δ¹⁵N, δ³⁴S, δ²H, δ¹⁸O), we estimated species-specific niche regions using a probabilistic, sample size-insensitive method implemented in the *nicheROVER* R package (Lysy et al. 2014) following the Bayesian niche region framework described by Swanson et al. (2015). For each species, 10,000 posterior draws were sampled from a Normal-Inverse-Wishart distribution, and niche size was defined as the hypervolume of the 95% Bayesian niche region. Directional pairwise niche overlap — the probability that an individual from one species falls within the niche region of another — was estimated using Monte Carlo simulations (10,000 steps) and summarised as mean overlap percentages with associated 95% credible intervals.

We began by exploring niche differentiation among ungulates through non-parametric Kruskal–Wallis tests on single isotope density distributions, followed by Dunn’s post-hoc tests in case of null hypothesis rejection (*FSA* R package; Ogle et al. 2015). To examine multivariate patterns in isotopic composition, consistent with predictions of niche differentiation along multiple ecological axes, we performed a Principal Component Analysis (PCA) on five isotope ratios (δ¹³C, δ¹⁵N, δ³⁴S, δ²H, δ¹⁸O), scaled to unit variance. We evaluated the proportion of variance explained by each axis and used loading scores and contribution metrics to interpret variable influence. To assess species-level differences in multivariate isotopic composition along principal components, we applied non-parametric Kruskal–Wallis tests to the PCA scores (i.e. sample positions in reduced isotopic space), followed by two-sided pairwise Dunn’s post-hoc comparisons in case of null hypothesis rejection. We visualised species clustering and overlap in isotopic space using individual score plots, and interpreted the underlying isotopic structure by plotting variable loadings (*factoextra* R package; Kassambara and Mundt 2020), providing insights into realised resource use.

All analyses were performed in R software (v.4.4.4; R Development Core Team 2022).

## Results

Based on the 95% Bayesian niche hypervolumes (Fig.S2), no strong differences in niche size were found among the three ungulate species (Fig.2). Red deer had a slightly broader niche (mean ± standard deviation: 8,962.24 ± 2,013.50) than roe deer (6,113.12 ± 2,445.29), with non-overlapping 50% credible intervals, but overlapping 90% credible intervals. Chamois niche size (7,516.45 ± 2,203.67) had overlapping 50% and 90% credible intervals with both roe deer and red deer (Fig.2).

**Figure 2.**
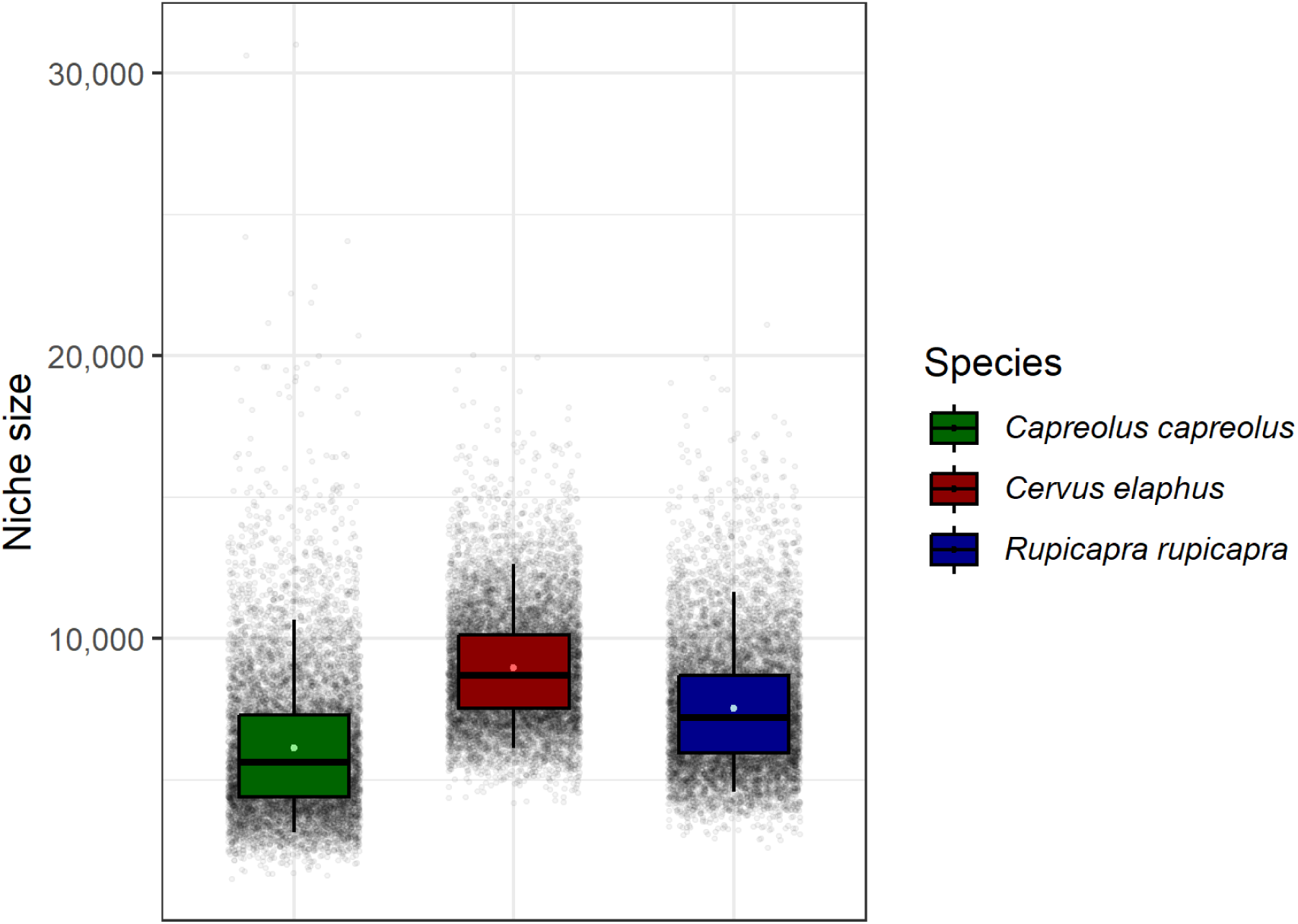
Niche size estimations, defined as the hypervolume of the 95% Bayesian niche region (relative values without units), for roe deer (left, green), red deer (centre, red), and chamois (right, blue). The plot shows the median niche size (horizontal line in the box), the mean niche size (light-coloured dot in the box), the 50% credible intervals (coloured box) and the 90% credible intervals (vertical lines). The background displays the 10,000 posterior niche size points.

Directional pairwise niche overlap analyses suggested that the mean overlap percentages between red deer, roe deer, and chamois never exceeded 40% (Fig.3). Specifically, the mean overlap probability of a roe deer being found within the red deer’s niche region was 30% (95% CI: 14.50 – 50.02), and within the chamois’ niche region was 25% (95% CI: 8.24 – 49.03); the mean overlap probability of a red deer being found within the roe deer’s niche region was 31% (95% CI: 14.79 – 51.20), and within the chamois’ niche region was 35% (95% CI: 16.72 – 57.09); and the mean overlap probability of a chamois being found within the roe deer’s niche region was 20% (95% CI: 5.80 – 43.33), and within the red deer’s niche region was 39% (95% CI: 20.56 – 58.98).

**Figure 3.**
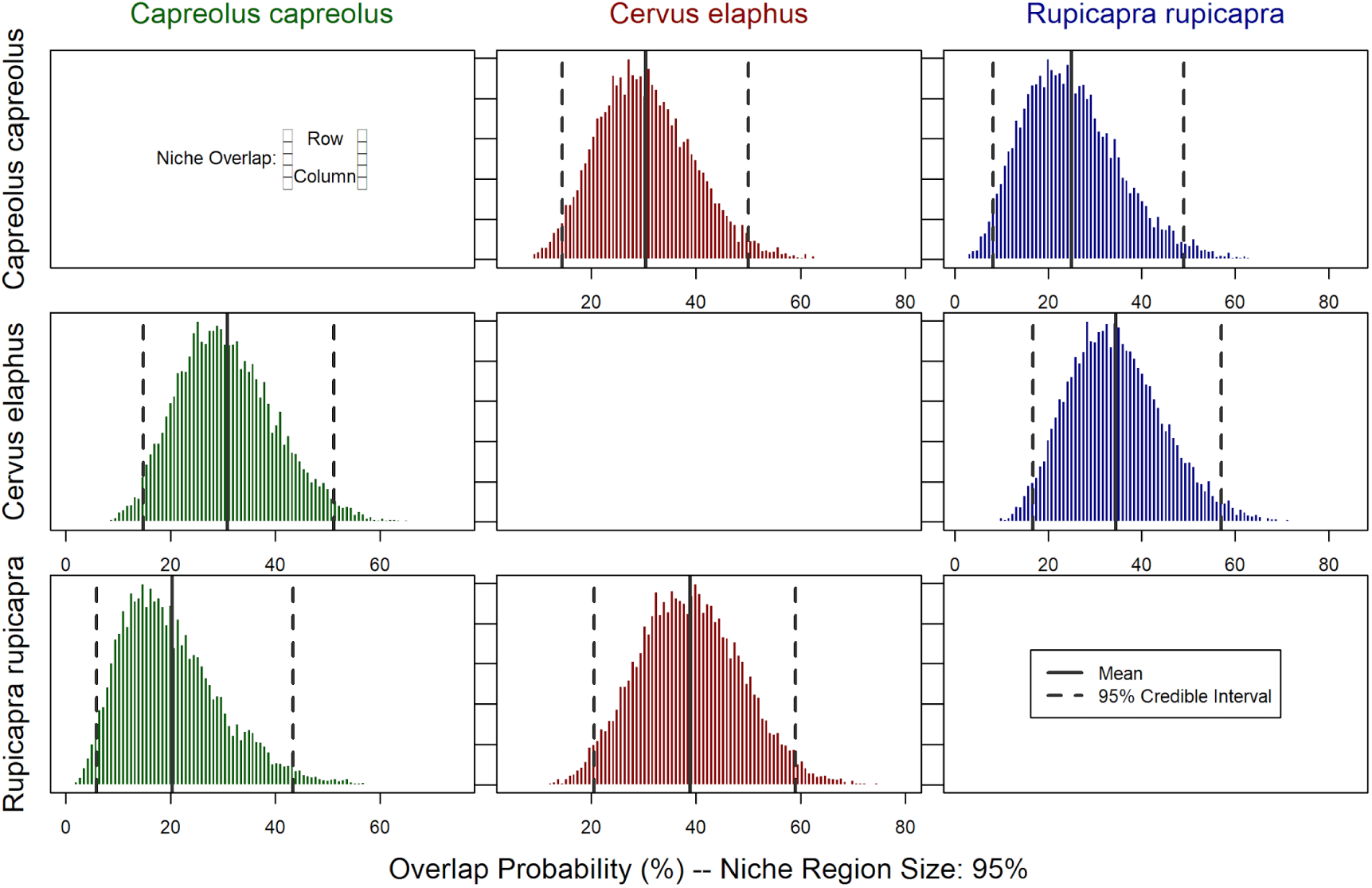
Posterior distribution of directional pairwise overlap probabilities between 95% Bayesian niche regions of roe deer, red deer, and chamois. These overlap probabilities represent the likelihood of any species (**row**; from top to bottom: roe deer, red deer, and chamois) being found within the niche region of any other species (**column**; from left to right: roe deer in green, red deer in red, chamois in blue). Mean overlap probabilities and 95% credible intervals are indicated by solid and dashed grey lines, respectively.

We tested interspecific differences in single isotope density distributions (Fig.4), using non-parametric Kruskal–Wallis tests and Dunn’s post-hoc tests. We found significant differences between species for all stable isotopes, except for δ^34^S (Table 3). Specifically, chamois had significantly lower δ^15^N and higher δ^18^O than both Cervids (Table 3), and red deer had significantly lower δ^2^H and δ^13^C than both roe deer and chamois (Table 3).

**Figure 4.**
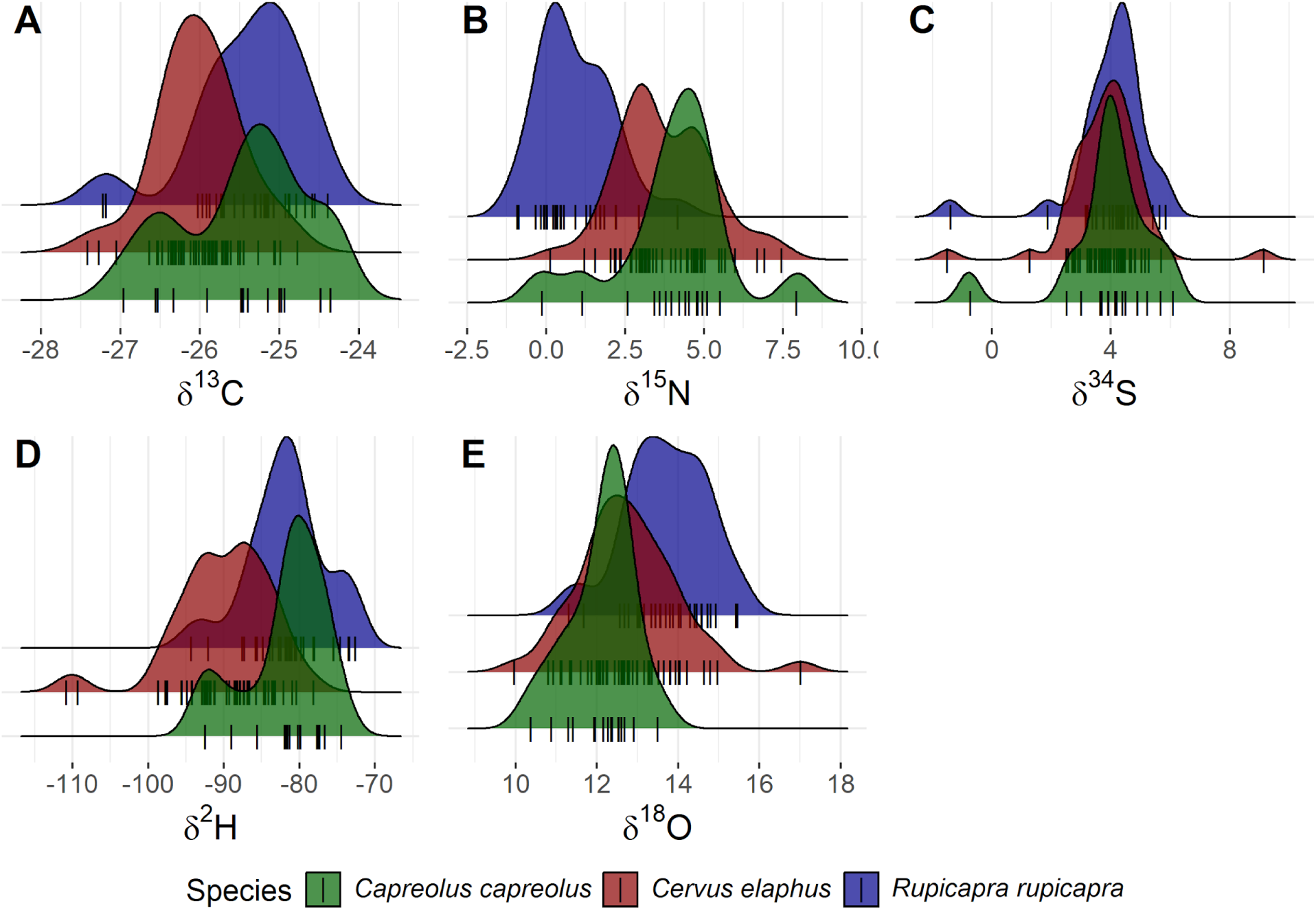
Density distributions of stable isotope values (δ^13^C, δ^15^N, δ^34^S, δ^2^H, and δ^18^O in **A**, **B**, **C**, **D**, and **E**, respectively), derived from untransformed individual-level measurements for roe deer (green, front), red deer (red, middle), and chamois (blue, back).

**Table 3.** Non-parametric Kruskal–Wallis tests on single isotope density distributions, followed by Dunn’s post-hoc tests in case of null hypothesis rejection.

|  | Kruskal-Wallis test |  | Post hoc multiple comparison Dunn’s test (two-sided, p-values adjusted with Bonferroni method) |  |  |  |  |  |
| --- | --- | --- | --- | --- | --- | --- | --- | --- |
|  |  |  | Roe deer vs Red deer |  | Roe deer vs Chamois |  | Red deer vs Chamois |  |
| | $\chi^2$ | $p$ | $z$ | $p_{adj}$ | $z$ | $p_{adj}$ | $z$ | $p_{adj}$ |
| $\delta^{13}\text{C}$ | 21.11 | < 0.001 | 2.88 | 0.012 | -0.54 | 1.000 | -4.27 | < 0.001 |
| $\delta^{15}\text{N}$ | 44.13 | < 0.001 | 0.65 | 1.000 | 5.19 | < 0.001 | 6.11 | < 0.001 |
| $\delta^{34}\text{S}$ | 1.78 | 0.412 | - | - | - | - | - | - |
| $\delta^2\text{H}$ | **35.71** | **< 0.001** | **4.35** | **< 0.001** | 0.11 | 1.000 | **-5.21** | **< 0.001** |
| $\delta^{18}\text{O}$ | **23.67** | **< 0.001** | -1.95 | 0.155 | **-4.55** | **< 0.001** | **-3.68** | **< 0.001** |

In the Principal Component Analysis, the first three components explained 77.49% of the total variance (PC1: 40.86%, PC2: 20.01%, PC3: 16.62%; Fig.S3), and were retained for further analysis and visualisation. **PC1** scores, primarily associated with high positive loadings of δ^2^H and δ^18^O, and a negative loading of δ^15^N (Table 4, Fig.5B,D), showed significant differences among species (Kruskal–Wallis test: χ² = 40.41, *p* < 0.001). Post hoc Dunn’s tests for PC1 indicated that chamois differed significantly from both red deer (Z = -6.35, *p_adj_* < 0.001) and roe deer (Z = -3.27, *p_adj_* = 0.003), while no significant difference was detected between red and roe deer (Z = 1.61, *p_adj_*= 0.322). These differences were driven by chamois’ relatively higher δ^2^H and δ^18^O, and lower δ^15^N values compared to Cervids (Fig.5). **PC2**, primarily associated with a high positive loading of δ^34^S (Table 4, Fig.5B), did not reveal significant interspecific differences (Kruskal–Wallis test: χ² = 4.93, *p* = 0.085; Fig.5A). Similarly, **PC3**, primarily associated with a high negative loading of δ^13^C (Table 4, Fig.5D), showed significant differences among species (Kruskal–Wallis test: χ² = 11.71, *p* = 0.003; Fig.5C). Post hoc Dunn’s tests for PC3 indicated that roe deer differed significantly from red deer (Z = -3.39, *p_adj_* = 0.002), while no significant difference was detected between roe deer and chamois (Z = -2.07, *p_adj_*= 0.117), nor between red deer and chamois (Z = 1.42, *p_adj_* = 0.465). These differences were driven by the relatively higher δ^13^C of roe deer compared to red deer (Fig.5).

**Figure 5.**
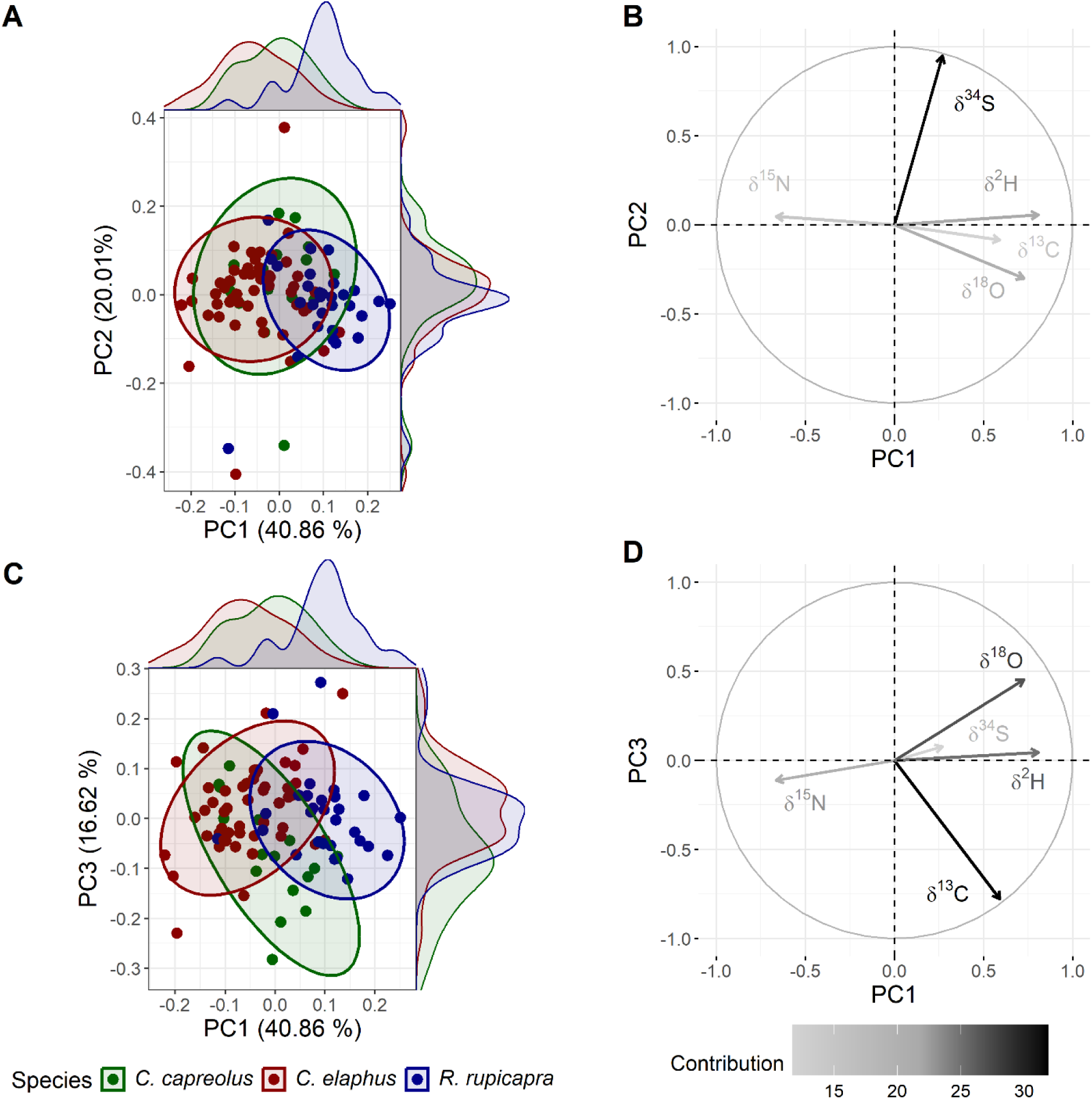
Ordination of the five stable isotope ratio variables (δ^13^C, δ^15^N, δ^34^S, δ^2^H, and δ^18^O) using PCA for roe deer (green), red deer (red), and chamois (blue). The scatterplots (**A,C**) show individual sample scores (dots) projected onto PC1 *vs* PC2 (**A**), and PC1 *vs* PC3 (**C**), with 95% ellipses drawn using multivariate t-distributions to show species-level niche dispersion. Marginal density plots (**A,C**; top and right sides) illustrate the distribution of principal component scores per species along each axis. The correlation circles for PC1 *vs* PC2 (**B**) and for PC1 *vs* PC3 (**D**) show how each isotope variable correlates with the principal components: arrow direction and length reflect correlation strength and orientation, while shading denotes variable contributions to the axes (from light grey to black). All panels use a fixed axis aspect ratio to ensure consistent scaling, facilitating visual comparisons across panels.

**Table 4.** Principal component contributions and loading scores. In bold are primary contributors (cumulating 70%). Underlined values are high loadings (> |0.4|).

|  | Contributions |  |  |  |  | Loading scores |  |  |  |  |
| --- | --- | --- | --- | --- | --- | --- | --- | --- | --- | --- |
|  | PC1 | PC2 | PC3 | PC4 | PC5 | PC1 | PC2 | PC3 | PC4 | PC5 |
| $\delta^{13}\text{C}$ | 17.12 | 0.73 | **72.84** | 0.13 | 9.19 | <u>0.41</u> | -0.09 | <u>-0.85</u> | 0.04 | 0.30 |
| $\delta^{15}\text{N}$ | **21.67** | 0.22 | 1.50 | **72.45** | 4.17 | <u>-0.47</u> | 0.05 | -0.12 | <u>0.85</u> | 0.20 |
| $\delta^{34}\text{S}$ | 3.54 | **89.70** | 0.71 | 0.00 | 6.05 | 0.19 | <u>0.95</u> | 0.08 | 0.00 | 0.25 |
| $\delta^{18}\text{O}$ | **25.76** | 9.05 | 24.71 | 4.96 | **35.52** | <u>0.51</u> | -0.30 | <u>0.50</u> | 0.22 | <u>0.60</u> |
| $\delta^2\text{H}$ | **31.91** | 0.31 | 0.24 | 22.47 | **45.07** | <u>0.56</u> | 0.06 | 0.05 | <u>0.47</u> | <u>-0.67</u> |

## Discussion

Understanding the functional structure of ecological communities and the underlying mechanisms for species coexistence within these communities is of critical importance to better predicting and mitigating consequences of rapid environmental change on natural communities and ecosystems (van der Putten et al. 2010; Sih et al. 2011; Ehrlén and Morris 2015; Gaüzère et al. 2018). Here, we tested isotopic niche-based hypotheses to assess the extent and axes of resource differentiation that facilitate coexistence among red deer, roe deer, and chamois sharing alpine summer ranges undergoing rapid environmental change (Passoni et al. 2024). In line with our predictions, our results indicate clear niche differentiation among these sympatric large herbivores, with limited isotopic niche overlap (<40%). Instead, we did not find significant differences in niche size among the three ungulates.

Building on this overall structure, species differed along individual isotopic axes in ways that partially matched our predictions (Table 1). As predicted for δ^13^C, in the univariate analyses, red deer showed lower values than chamois, consistent with greater use of forested habitats, while roe deer were intermediate. In contrast, isotopes typically associated with elevation (δ^2^H, δ^18^O) did not follow the expected altitudinal patterns: red deer had lower δ^2^H values than chamois and roe deer, and chamois had unexpectedly high δ^18^O values, possibly suggesting species-specific differences in water sourcing (Hobson et al. 1999; Farquhar and Gan 2003). Dietary differentiation contradicted the predicted pattern for δ^15^N. While red deer exhibited higher δ^15^N as expected, chamois showed significantly lower δ^15^N values than both Cervids, suggesting that the realised summer diet of chamois is of higher quality than that of red deer and roe deer. Fine-scale diet composition, protein content and nutrient routing (Short et al. 1974; Delwiche et al. 1979; Mattson 1980; Ambrose and DeNiro 1986; Robbins et al. 1987; Fantle et al. 1999; Vanderklift and Ponsard 2003; Hobson 2008) appear to override the simplistic browser-grazer continuum (Codron et al. 2007), highlighting that realised diet quality can vary substantially even within broad feeding categories. As predicted for δ^34^S, no significant interspecific differences were observed, indicating shared use of areas at the geological scale. Multivariate analyses provided complementary support for these patterns: chamois separated from Cervids primarily through δ^15^N, δ^2^H, and δ^18^O, whereas Cervids separated primarily through δ^13^C, reinforcing the conclusion that isotopic profiles reflect functional differentiation in diet quality, water sourcing, and canopy cover use. Together, these results highlight how stable isotope variation captures niche differentiation that likely underpins coexistence in this Alpine ungulate assemblage.

### Diet quality, water uptake source, and canopy cover use as main factors of differentiation

#### δ^15^N: diet quality

Previous studies have demonstrated a positive relationship between dietary C:N ratio — a reliable indicator of dietary protein content, fiber-to-protein composition, and diet quality (Hobson 2008) — and nitrogen isotope discrimination (Cormie and Schwarcz 1996; Fantle et al. 1999; Vanderklift and Ponsard 2003). We expected that δ^15^N signatures would therefore reflect ungulates’ positioning on the browser-grazer continuum (Table 1), specifically with browsers (or concentrate selectors like roe deer; Hofmann 1989) exhibiting higher δ^15^N values than intermediate feeders (chamois and red deer; Hofmann 1989). Instead, our results showed lower δ^15^N values in chamois compared to red deer and roe deer, a pattern consistently observed in single isotope comparisons (Fig.4B) and in the principal component analysis (Fig.5). This finding suggests that fine-scale diet composition and nutrient routing may override the somewhat simplistic browser-grazer continuum (Codron et al. 2007).

Specifically, consumption of highly palatable and protein-rich food items with low C:N ratios (whether belonging to grasses, forbs, shrubs, trees or other functional plant groups) lead to lower δ^15^N values in consumer tissues. Hence, lower δ^15^N in chamois may indicate a diet more largely based on such palatable food items, confirming the major role of biomechanical plant traits in chamois diet selection (Bison 2015), as well as for their high consumption of forbs (20.98% of chamois diet *vs* 4.11% of red deer diet; Redjadj et al. 2014). Leguminous nitrogen fixing plants are also typically ^15^N-depleted (Delwiche et al. 1979; Ambrose and DeNiro 1986) and may thus be selected by chamois, confirming findings from the French Prealps where chamois consumed about five and fourteen times more leguminous nitrogen-fixing plants than red deer and roe deer, respectively (Redjadj et al. 2014). In contrast, the higher δ^15^N in red deer tissues may indicate a higher ingestion of grasses or fibrous forage (Hofmann 1989; Codron et al. 2007) as predicted (Table 1), but also more fruits and seeds with high C:N composition, such as acorns or apples (Gebert and Verheyden-Tixier 2001; Sun et al. 2012; Redjadj et al. 2014). Consumption of lignified tissue and tannins can also reduce overall diet quality (higher C:N) and nitrogen availability (Short et al. 1974; Mattson 1980; Robbins et al. 1987), which may contribute to the elevated δ^15^N values in roe deer. Indeed, as their rumen is less adapted to digest fiber (Clauss et al. 2010), roe deer typically browse on forbs, shrubs and trees (79.08% of diet, of which 48.18% on evergreen shrubs in the French Prealps; Redjadj et al. 2014), thereby ingesting significant quantities of lignified tissue and tannins (Tixier et al. 1997; Verheyden-Tixier and Duncan 2000; Clauss et al. 2003).

Additional factors may also contribute to the observed δ^15^N patterns, including interspecific differences in the selection of plant phenological stages (Aikens et al. 2017; Shamon et al. 2022) or plant organs (e.g., leaves *vs* stems; Gwynne and Bell 1968); these may also be linked to differences in palatability, protein content, and C:N ratio. The observed δ^15^N patterns may further be shaped by interspecific differences in digestive function, including anatomical adaptations (Clauss et al. 2008) and gut microbiota composition (Qin et al. 2020; Zhang et al. 2025). Furthermore, δ^15^N values in consumer tissues can be influenced by complex, interdependent mechanisms involving not only diet composition but also abiotic factors such as precipitation (Heaton et al. 1986; Ambrose 1991; Cormie and Schwarcz 1996). Disentangling the influence of these factors on the isotopic patterns of sympatric ungulates warrants further investigation.

#### δ^2^H, δ^18^O: water sourcing (and fine-scale topographic complexity)

The observed patterns in δ^2^H and δ^18^O among ungulates did not align clearly with elevational predictions (Table 1), which would have reflected the altitudinal depletion of heavy isotopes in precipitation water (Dansgaard 1964). Instead, we found that chamois had higher δ^18^O values than both Cervid species (Fig.4E, Fig.5A-B), while δ^2^H values were lowest in red deer, and similar between chamois and roe deer (Fig.4D, Fig.5C-D). These results suggest that elevation alone does not explain the observed isotopic variation.

Instead, the differences in δ^18^O we found were similar to results found in the French Prealps (Merceron et al. 2021), and may reflect variation in the sources of water uptake: chamois may derive a larger proportion of their water from vegetation and metabolic water — both enriched in ^18^O and ^2^H relative to environmental water (Hobson et al. 1999; Farquhar and Gan 2003) — whereas red and roe deer may rely more on free water from streams, snowmelt, or dew. (There is a general scarcity of literature on drinking behaviour, especially for chamois, but see (Wallach et al. 2007) for roe deer). It is worth noting that, although chamois may drink less, we did not observe isotopic evidence of water stress or dehydration, which would typically manifest as elevated δ^15^N values due to changes in nitrogen metabolism under limited water availability (Ambrose and DeNiro 1986; Heaton et al. 1986; Ambrose 1991; Sponheimer et al. 2003). Beyond differences in water uptake, the observed interspecific variation in δ^2^H and δ^18^O values may also reflect the effects of fine-scale topographic complexity (Loader et al. 2016). In rugged and heterogeneous alpine terrain, microclimatic conditions vary locally with slope, aspect, and exposure, potentially obscuring clear elevational isotopic patterns. This spatial heterogeneity complicates the interpretation of δ^2^H and δ^18^O values in herbivore tissues and highlights the need for more targeted, time-resolved studies, particularly at fine spatial scales, to refine their interpretation in large herbivore tissues (Burnik Šturm et al. 2017).

#### δ^13^C: habitat use

Ground vegetation in densely forested areas are exposed to lower light intensity (affecting stomatal conductance and Rubisco carboxylation; Farquhar et al. 1982; Ehleringer et al. 1986; Broadmeadow et al. 1992) and higher CO_2_ concentrations (^13^C-depleted CO_2_ from biomass degradation and respiration; Vogel 1978; Da Silveira et al. 1989) than vegetation in open areas, leading to lower δ^13^C signatures in forest ground vegetation (van der Merwe and Medina 1991; Heaton 1999; Bonafini et al. 2013; Table 1). In pure C_3_ ecosystems, such as the Alps, relative resource use from forested *vs* open environments by herbivores may be inferred in this way (Drucker et al. 2008; Drucker and Bocherens 2009; Drucker et al. 2010).

At the population level, red deer exhibited significantly lower δ^13^C values than chamois, suggesting a greater reliance on forest-derived resources relative to the alpine grasslands and shrubland habitats used by chamois (von Elsner-Shack 1985; Nesti et al. 2010). These results should not be interpreted as excluding the use of open habitats by red deer, but rather as indicative of a greater relative reliance on resources from forested areas within the sampled population. A similar contrast in δ^13^C values between these species was also reported from tooth enamel in the French Prealps (Merceron et al. 2021). Roe deer, a forest-dwelling browser, was also expected to show lower δ^13^C values consistent with forest use. However, instead, they displayed significantly higher δ^13^C values than red deer (Fig.4A, 5A), possibly reflecting their greater reliance on resources from ecotonal habitats (Andersen et al. 1998), where light penetration and air ventilation are greater than inside dense forest cover. Moreover, a vertical δ^13^C gradient occurs within forests, with δ^13^C values increasing with height above the ground (Vogel 1978; E. Medina and Minchin 1980; Ernesto Medina et al. 1986). This pattern may further contribute to roe deer isotopic signatures, as they are concentrate selectors, often feeding on higher shrubs and twigs.

#### δ^34^S: local geology

Sulphur isotopes in herbivore tissues are typically influenced by underlying geology and soil composition (Richards et al. 2003; Nehlich 2015; Table 1). Although a significant portion of the total variance among samples was linked to δ^34^S (PC2 primarily structured by δ^34^S, 20.01% of total variance; Table 4, Fig.5A), this variance did not reflect interspecific differences (Fig.4C, Fig.5A). Instead, our findings indicate variation in geochemical conditions among individuals, but no differentiation at the species level, pointing to spatial overlap at the considered scale. In contrast to the marked interspecific differences observed for other isotopes, the uniformity in δ^34^S highlights the value of considering and integrating multiple bio-elements, each bringing forth different axes of ecological relevance.

#### Other factors influencing isotopic signatures

Physiological and metabolic states, including growth rate and nutritional stress, as well as abiotic conditions such as temperature, soil moisture or atmospheric humidity, may further modulate isotopic incorporation of carbon, sulphur, nitrogen, hydrogen and oxygen (Hobson et al. 1993; Fantle et al. 1999; Drucker et al. 2008; Caut et al. 2009; Shipley and Matich 2020; Kabalika et al. 2024), highlighting the necessity for future studies to integrate dietary, physiological, and environmental data to isolate the specific drivers of isotopic variation. Disentangling these processes is essential for moving beyond a descriptive interpretation of isotopic niche differentiation toward a mechanistic understanding of underlying ecological niche segregation among sympatric ungulates.

### Isotopic differentiation despite overlaps as a mechanism for coexistence

To explain the coexistence of closely related species, two main niche-based mechanisms have been proposed: niche filtering and niche partitioning. Niche filtering suggests that environmental conditions select for species with similar, environment-adapted traits (Keddy 1992; Zobel 1997), while the limiting similarity hypothesis posits that competitive exclusion limits coexistence among too-similar species, leading to niche partitioning (Macarthur and Levins 1967; Chesson 2000a; Chase and Leibold 2003; Barabás et al. 2018).

In this study, mean pairwise isotopic niche overlaps among red deer, roe deer and chamois ranged from 20% to 39%, with upper 95% CI limits never exceeding 59% (Fig.3). These values fall below commonly cited thresholds for biologically meaningful overlap (typically 60–70%; Zaret and Rand 1971; Wallace 1981; Wathne et al. 2000; Guzzo et al. 2013), suggesting substantial niche segregation despite some degree of shared resource use. Consistent with this, our results show clear isotopic segregation along certain niche dimensions, but convergence along others. For example, red deer and roe deer were only significantly differentiated by δ^13^C and δ^2^H, markers associated with habitat openness and abiotic/microclimatic conditions (see previous section), while they overlapped in δ^34^S, δ^15^N, and δ^18^O, reflecting shared spatial ranges at the geological and limnological scales, similar diet quality, and comparable water sourcing. Chamois and red deer overlapped in δ^34^S, likely reflecting shared spatial ranges at the geological and limnological scales, but were clearly separated along δ^2^H, δ^18^O, δ^15^N, and δ^13^C, suggesting differences in abiotic/microclimate exposure, water sourcing, diet quality and habitat use (see previous section). Chamois and roe deer only differed in δ^15^N and δ^18^O, suggesting differences in feeding and drinking behaviour (see previous section).

Together, these findings suggest that both niche partitioning and niche filtering operate concurrently within this Alpine ungulate community — a pattern observed in other systems, where differentiation along certain niche dimensions may facilitate filtering along others (Spasojevic and Suding 2012). For example, coexisting foliage-dwelling spiders (*Philodromus spp.*) were found to strongly overlap along habitat and spatial niche dimensions but to partition along their functional niche dimensions (involving prey size, body size and aggressiveness towards prey; Michalko and Pekár 2015). Similarly, filtering and differentiation jointly determined species abundance in grassland communities by acting on different niche dimensions, with niche overlap being higher than expected at random for traits like plant height, leaf and stem dry matter content, and leaf C:N ratio, whereas it was lower than expected for specific leaf area (Mason et al. 2011; Maire et al. 2012). These axis-specific patterns of segregation support the idea that distinct ecological dimensions can be governed by different coexistence mechanisms, with partitioning along traits such as diet or water acquisition potentially facilitating filtering along others, like elevation or habitat use. Rather than representing a contradiction, such asymmetry aligns with a multidimensional view of community assembly (Schoener 1974), in which different niche dimensions are structured by distinct ecological processes (Kraft, Godoy, et al. 2015; Kraft, Adler, et al. 2015; Cadotte and Tucker 2017).

### Beyond niche size and differentiation: integrating approaches to infer competition

Previous long-term demographic studies in Stelvio National Park have reported negative correlations between red deer density and chamois demographic parameters, such as growth rate and survival (Corlatti et al. 2019; Donini et al. 2021), which have been interpreted as indicative of interspecific competition, although the underlying mechanisms remained untested. At the same time, these and other studies across the Alps emphasise the strong influence of climate and habitat change (Garel et al. 2011; Rughetti and Festa-Bianchet 2012; Willisch et al. 2013) in shaping chamois population dynamics. Together, these findings point to a multifaceted ecological context in which competition and environmental change may interact, underscoring the complexity of disentangling the drivers of community structure in natural systems under anthropogenic influence.

Our study complements these population-level investigations by providing a functional ecological lens on realised resource use. Using stable isotope analyses, we examined multidimensional niche overlap and configuration to infer the underlying functional differences in resource acquisition strategies and environmental interactions within a sympatric ungulate community. Our results reveal clear isotopic segregation among red deer, chamois, and roe deer, likely reflecting differences in feeding and drinking behaviour, habitat use, and fine-scale topographic exposure. Importantly, we did not observe niche partitioning *sensu stricto* — a term that implies a process resulting from past or ongoing competition (Schoener 1974; Colwell and Fuentes 1975; Walter 1991; Matich et al. 2021). Instead, we interpret our findings as niche differentiation or segregation, which can result from multiple, potentially concurrent ecological processes (Michalko and Pekár 2015; Kraft, Adler, et al. 2015), including within-species individual niche variation which may further shape community dynamics (Bolnick et al. 2003, 2007; Lehmann et al. 2015).

Isotopic niche segregation reflects differences in realised resource use (Lehmann et al. 2015), and can serve as a valuable first step in assessing the potential for resource competition. While it is not direct evidence for (nor against) competition, it provides insight into patterns of resource segregation. Indeed, demonstrating resource competition requires not only identifying a shared resource, but also proving that it is limiting, and documenting overlap during the time of limitation (Walter 1991). Because they integrate over weeks to months (Hette-Tronquart 2019), stable isotope signatures in hair lack the temporal resolution to detect behavioural or interference competition, and cannot directly capture mechanisms of interaction or exclusion (Matich et al. 2021). Nonetheless, they provide an important basis for understanding species coexistence by indicating niche dimensions where overlap occurs and where further investigation is needed to formally assess competition.

To rigorously test for interspecific competition among red deer, roe deer and chamois, isotopic analyses could be integrated with complementary approaches that offer finer taxonomic, temporal, and spatial resolution. These include diet composition analyses, behavioural observations or spatial tracking (Shipley and Matich 2020; Merceron et al. 2021; Petalas et al. 2024), compound-specific isotope analyses (e.g. amino acid δ^15^N; van Oordt et al. 2024), or experimental and quasi-experimental study designs of community composition (Codron et al. 2011).

All studied ungulates exhibited similar isotopic niche sizes (Fig. 2), suggesting comparable breadth in resource use among hypothesised generalist (red and roe deer) and specialist (chamois) species. Thus, we found no isotopic evidence for differential niche breadth or isotopic generalism, traits that can be associated with resilience and plasticity under environmental change (Clavel et al. 2011; Sunday et al. 2015; Hurtado et al. 2024). However, such differences may still occur at temporal, spatial, or behavioural scales not captured by hair stable isotopes.

## Conclusions

Our results provide important insights into the functional structure of an Alpine ungulate community and the underlying mechanisms for coexistence. Future research should integrate isotopic analyses with complementary behavioural, trophic, demographic, or historical datasets to better disentangle the mechanisms underlying resource use and potential competition. For example, combining stable isotope data with analyses of museum specimens could reveal temporal shifts in niche use and potential signatures of past or evolving competition, while integrating isotope data with ruminant gut microbiota could shed light on the role of digestive capacities in shaping differential diet composition or nutrient uptake. Such integrative approaches will not only strengthen our mechanistic understanding of community dynamics, but also improve our ability to predict how communities respond to environmental and ecosystemic change.

## Statement of authorship

Conceptualisation (study design): CV, FC, LB. Conceptualisation (overarching project design): CV, LC, HCH, LP, FC. Data Curation: CV, AR, AT, LZ. Methodology: CV, LB. Formal Analysis: CV. Assessment of Model Outputs: CV, FC, LB. Investigation (field): CV, MN, LL. Investigation (laboratory): CV, GG, KW, AT, LZ. Software: CV. Validation (laboratory): AT, LZ. Validation (data analysis): CV. Visualisation: CV. Writing – Original Draft: CV. Writing – Review and Editing: CV, AR, LL, LC, HCH, FC, LB. Funding Acquisition: HCH, LP, FC, LB. Project Administration: LP, LC, FC, LB. Resources: FC, LB. Supervision: HCH, LP, FC, LB.

## Supplementary Materials

### Data S1. Stable isotope ratio determination

#### Sample preparation

Hair samples were cleaned and defatted via three successive 1-minute sonicated solvent baths at 40°C (diethyl ether:methanol 2:1). Samples were then placed in the oven at 50°C for at least one hour, to ensure evaporation of the solvent.

For δ^13^C, δ^15^N, and δ^34^S measurements, 0.450 mg [0.400–0.500] were weighed and placed in tin capsules, including 3 technical replicates. For δ^2^H and δ^18^O measurements, 0.250 mg [0.225–0.275] were weighed and placed in silver capsules, including 2 technical replicates.

#### International reference materials and working in-house standards

Values of δ^13^C, δ^15^N and δ^34^S were simultaneously determined using a Vario Isotope Cube isotope ratio mass spectrometer (Elemental, Germany). Stable isotope ratios were calculated against international reference standards: USGS 88 (United States Geological Survey, marine collagen from wild caught fish; δ^13^C = -16.06‰; δ^15^N = 16.96‰; δ^34^S = 17.1‰), USGS 90 (millet flour from Tuscany; δ^13^C = -13.75‰; δ^15^N = 8.84‰; δ^34^S = -15.14‰) and one working in-house standard (wheat; δ^13^C = -26.01‰; δ^15^N = 4.61‰; δ^34^S = 1.05‰), which was itself calibrated against international reference materials using multi-point normalisation: fuel oil NBS-22 (IAEA International Atomic Energy Agency, Vienna, Austria; δ^13^C = -30.03‰), sugar IAEACH-6 (δ^13^C = -10.45‰), L-glutamic acid USGS 40 (δ^13^C = -26.39‰; δ^15^N = -4.5‰), hair USGS 42 (δ^15^N = 8.05‰), rice flour USGS 91 (δ^34^S = -20.85‰), barium sulfate IAEA SO 5 (δ^34^S = 0.5‰), and barium sulfate NBS-127 (δ^34^S = 20.3‰).

Values of δ^2^H and δ^18^O were determined through pyrolysis combustion using a TC/EA (Thermo Finnigan, Bremen, Germany) interfaced with a Delta Plus XP (Thermo Finnigan, Bremen, Germany) continuous-flow isotope ratio mass spectrometer. Pyrolysis was carried out at 1450 °C in a glassy carbon column. The helium carrier flow was 110 ml/min, the GC column was a 1.2 m long molecular sieve 5A, at 110 °C. Stable isotope ratios were calculated against the reference materials CBS (Caribou Hoof Standard; δ^2^H = -157‰; δ¹⁸O = 3.8‰) and KHS (Kudu Horn Standard; δ^2^H = -35.3 ‰; δ¹⁸O = 20.3 ‰).

### Data S2. Bayesian niche region estimation

**Figure S.2.**
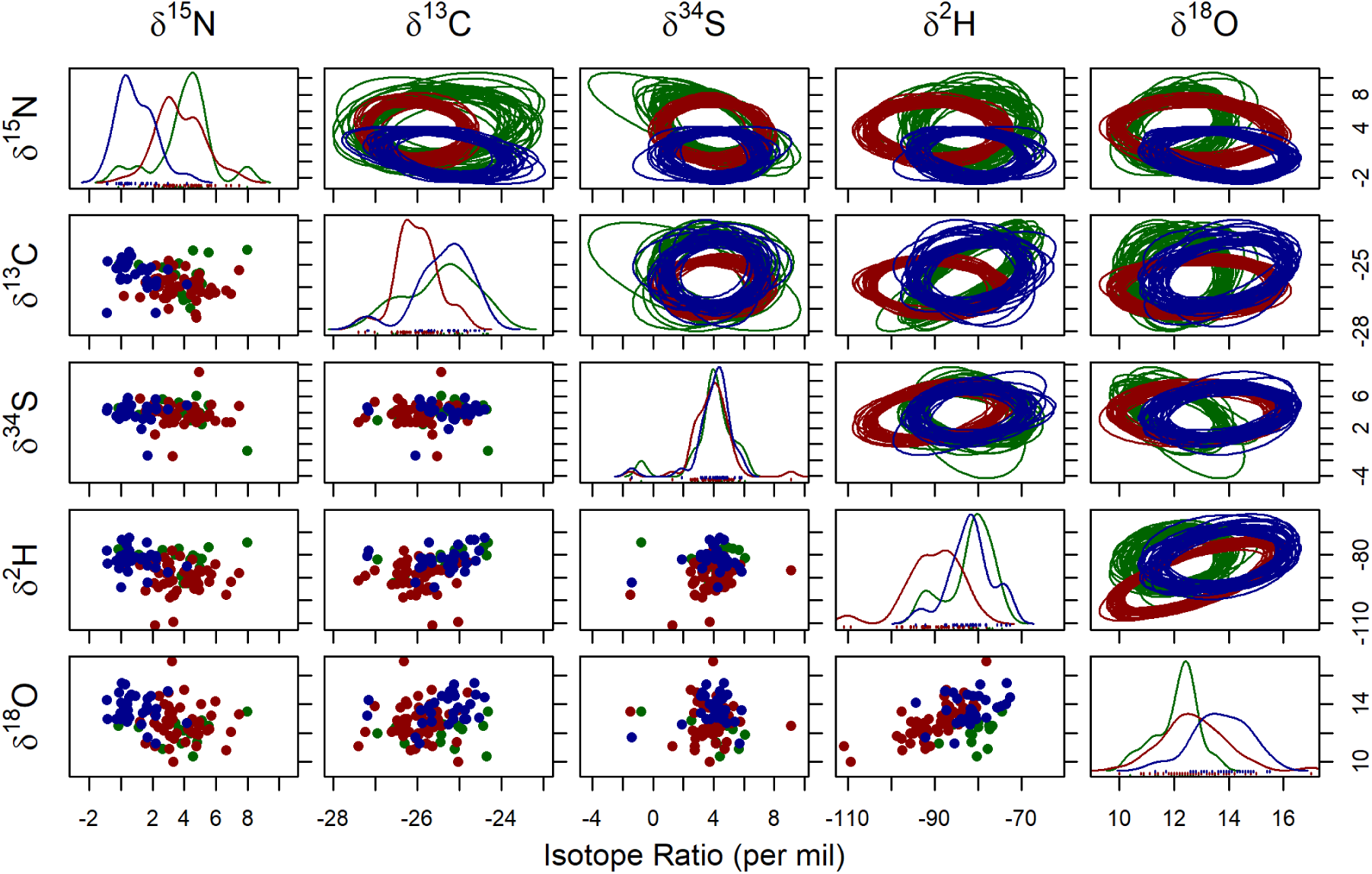
Using a Bayesian framework, 30 random elliptical projections (elliptical plots, top right) were drawn for red deer (red), roe deer (green), and chamois (blue). Also displayed are one-dimensional density plots (lines, center diagonale) and two-dimensional scatterplots (points, bottom left).

### Data S3. Principal Component Analysis

**Figure S.3.**
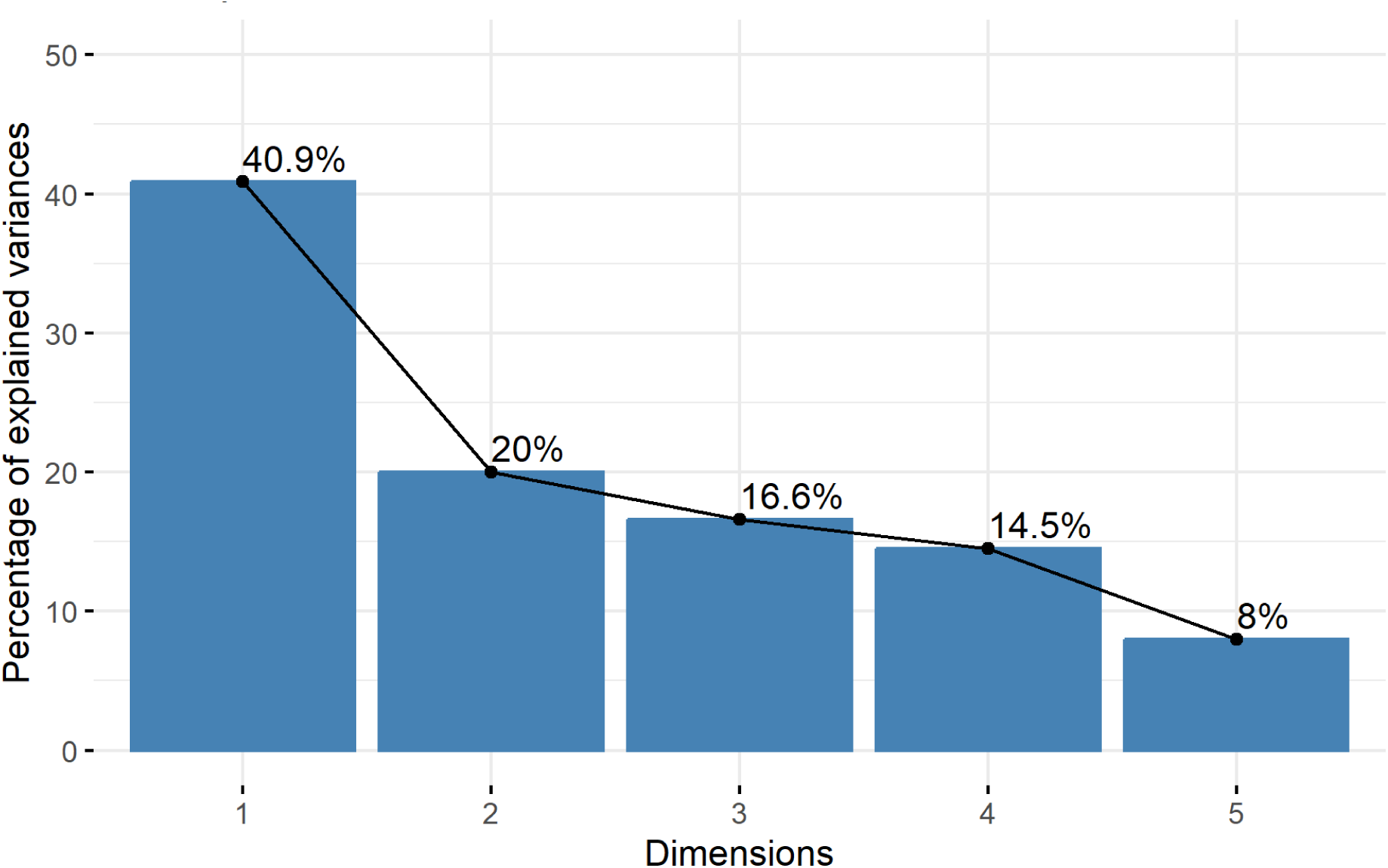
Scree plot showing the proportion of variance explained by each principal component (or dimension) in the PCA.

